## Supplemental file for "Structural and mechanistic insights into the MCM8/9 helicase complex"

### Supplementary figures

Figure S1

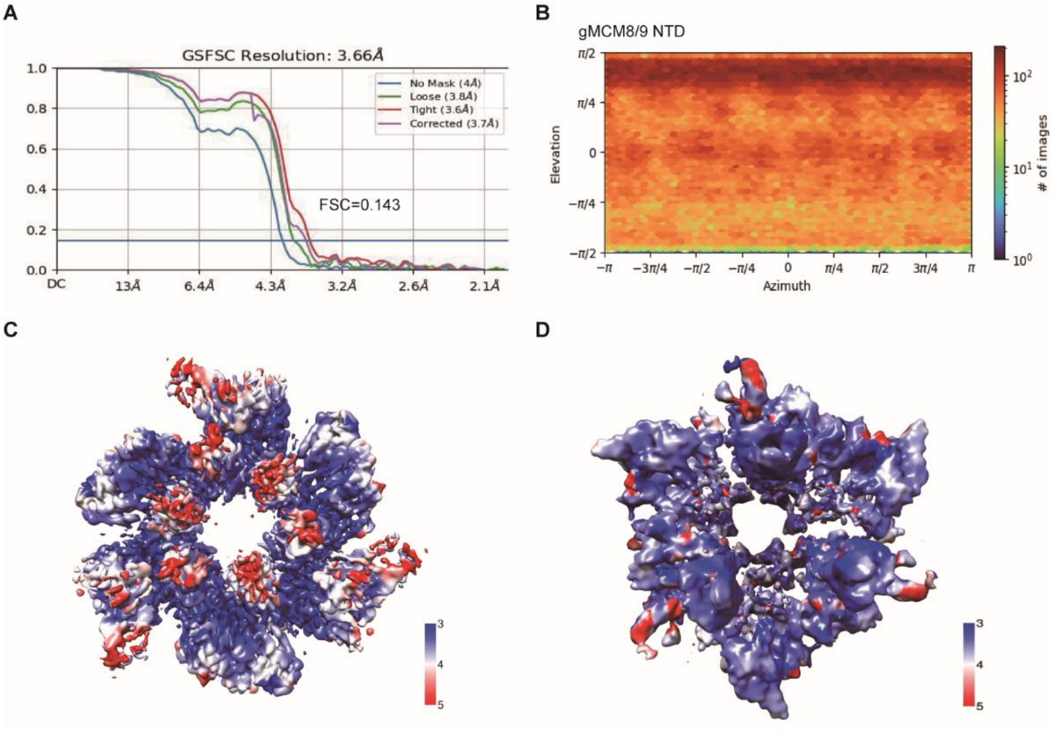


**Figure S1. Resolution evaluation of the gMCM8/9 NTD and CTD.** (**A**) Gold standard Fourier shell correlation (FSC) curve for the 3D refinement of the overall structure of the gMCM8/9 NTD. (**B**) Angular distribution of the particles used for the final reconstructions. (**C**) and (**D**) Local resolution distributions of gMCM8/9 NTD and gMCM8/9 CTD.

Figure S2

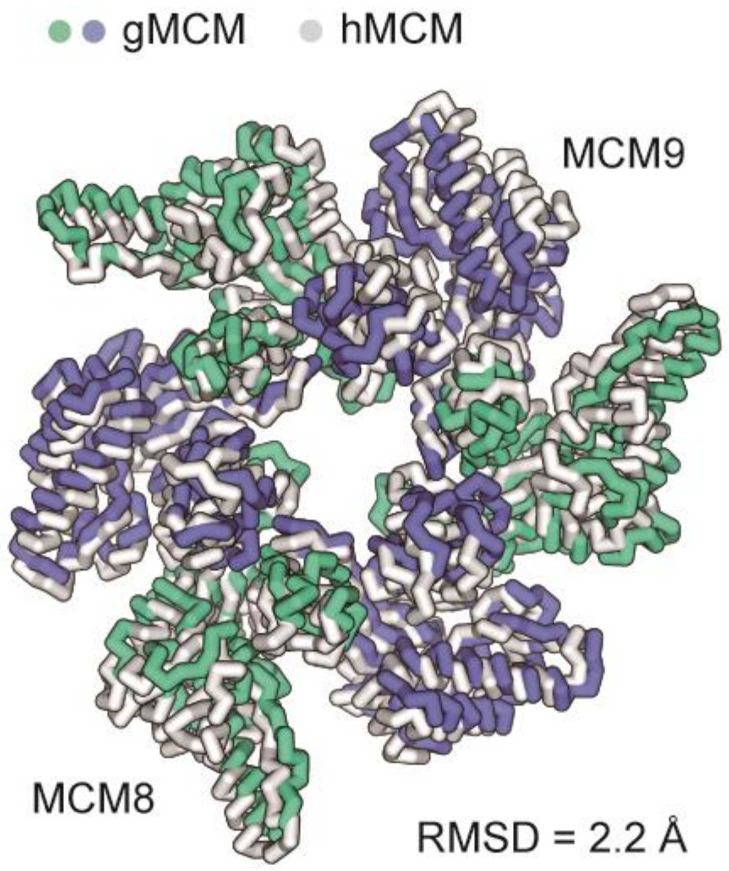


**Figure S2. Structural superposition of the NTDs of gMCM8/9 and hMCM8/9.** The overall root mean square deviation (RMSD) (2.2 Å) was listed below the structures. The structures were shown as ribbons. gMCM8/9, greencyan and slate; hMCM8/9, grey.

Figure S3

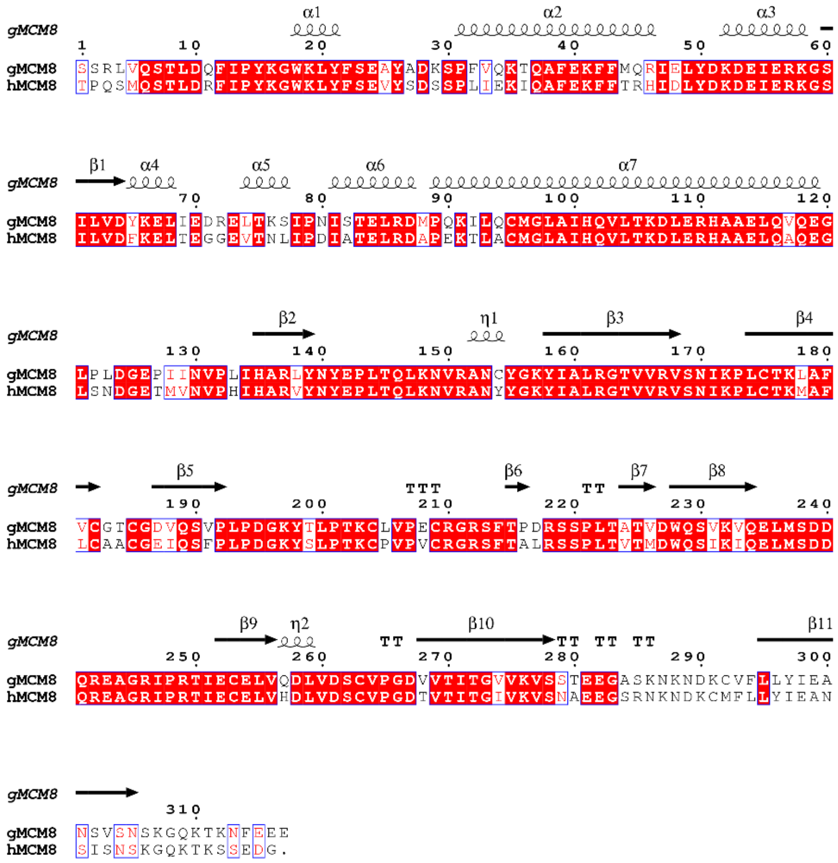

**Figure S3. Sequence alignments of NTDs of gMCM8 and hMCM8**. The secondary structures were shown above the sequences and the highly conserved residues were highlighted as red color. h, human; g, chicken (*Gallus gallus*).

Figure S4

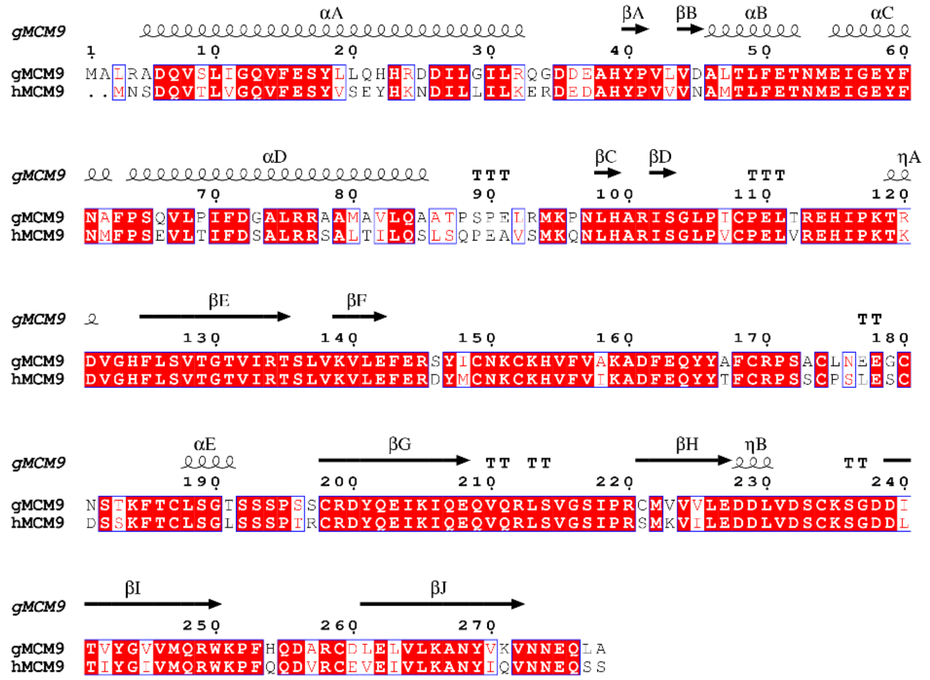

**Figure S4. Sequence alignments of NTDs of gMCM9 and hMCM9.** The secondary structures were shown above the sequences and the highly conserved residues were highlighted as red color. h, human; g, chicken (*Gallus gallus*).

Figure S5

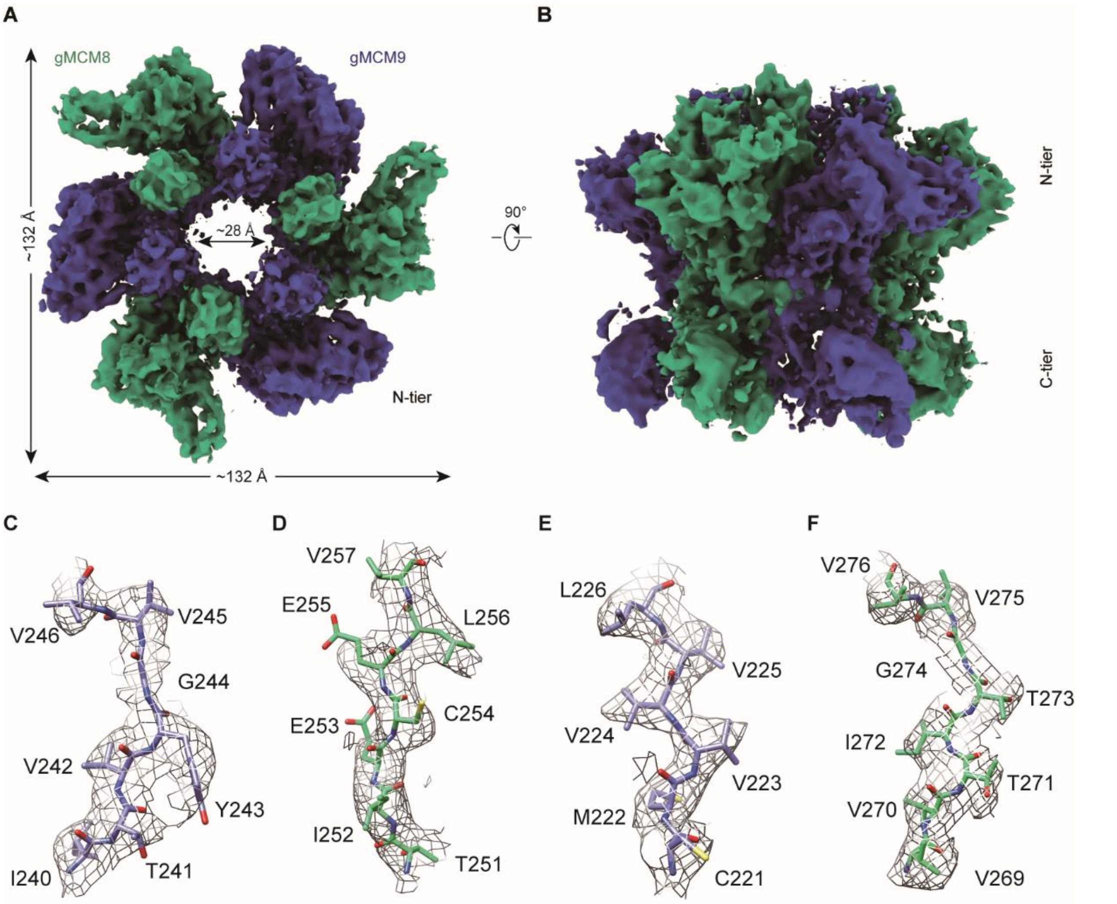


**Figure S5. The cryo-EM structure of gMCM8/9. (A and B)** Reconstructed cryo-EM map of gMCM8/9. The diameter of the inner channel of MCM8/9 was measured at ~28 Å. **(C-F)** Representative regions of the cryo-EM structure of gMCM8/9 NTD are shown based on their density map. C, chain A (MCM9); D, chain B (MCM8); E, chain C (MCM9); F, chain D (MCM8).

Figure S6

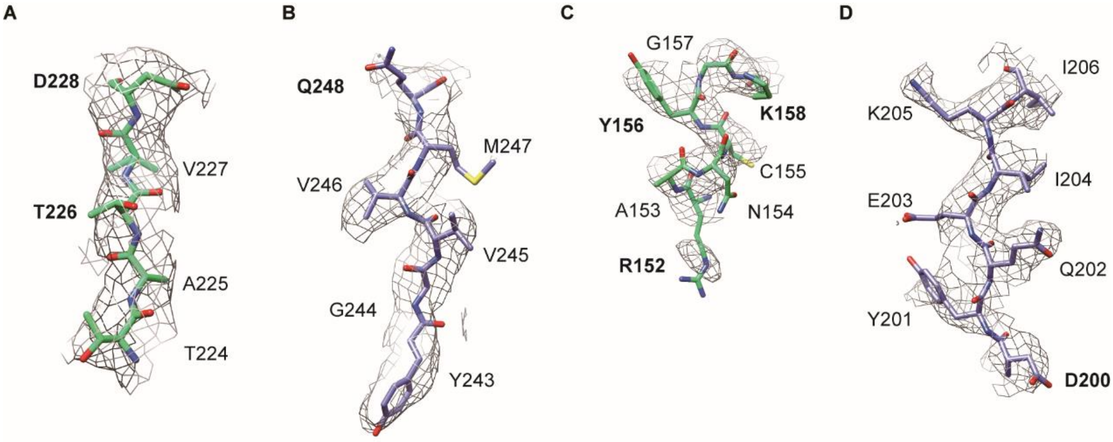


**Figure S6. Representative regions of the cryo-EM structure of gMCM8/9 NTD.** (**A and B**), the region mediated hydrophobic interaction in figure 2B. A (MCM8), B (MCM9). (**C and D**), the region mediated hydrophobic interaction in figure 2C. C (MCM8), D (MCM9). The key residues were in bold.

Figure S7

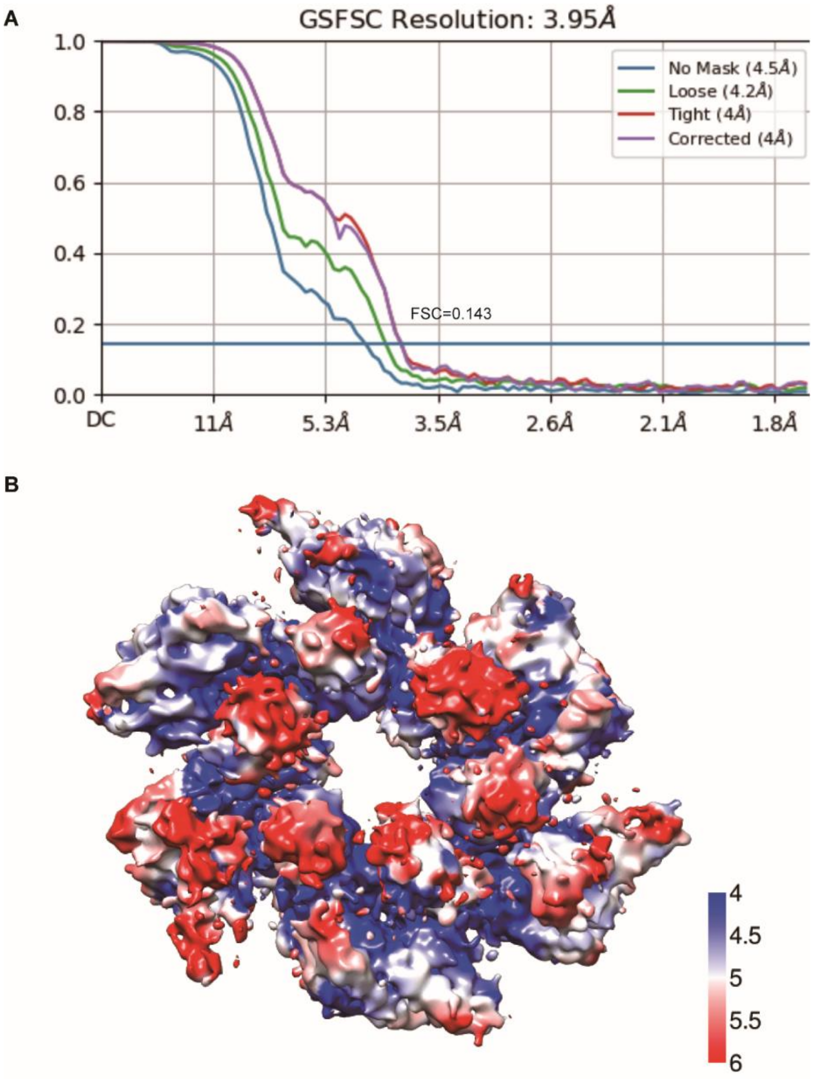


**Figure S7. Resolution evaluation of the hMCM8/9 NTD Conformation II.** (**A**) Gold standard Fourier shell correlation (FSC) curve for the 3D refinement of the overall structure of the hMCM8/9 NTD. (**B**) Local resolution distributions of hMCM8/9 NTD.

Figure S8

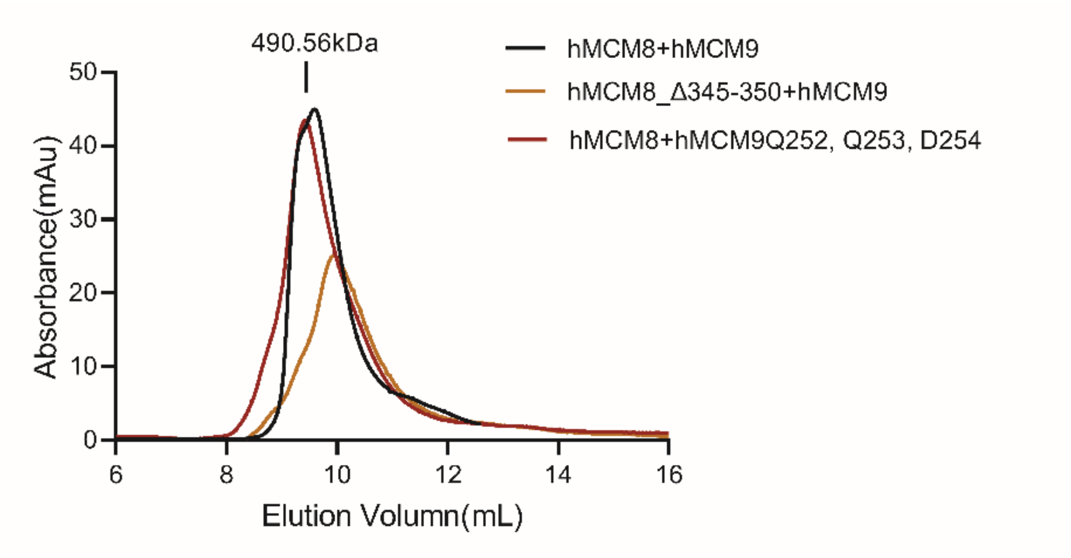


**Figure S8. SEC profiles of WT and OB-hps mutants of MCM8/9 complex.**

Figure S9

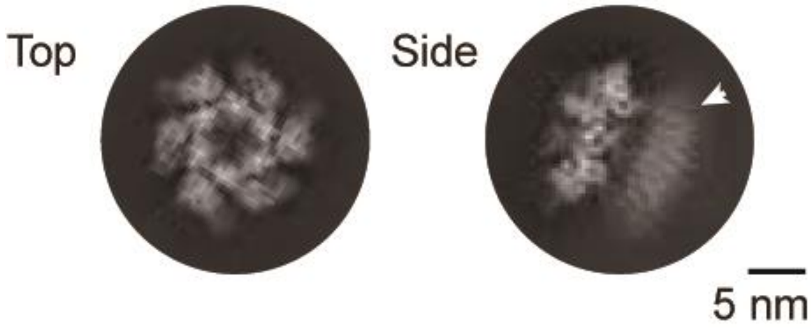


**Figure S9. Selected 2D classes of gMCM8/9 showing its typical top and side views.** The white triangle marks the blurry C-tier ring of gMCM8/9.

Figure S10

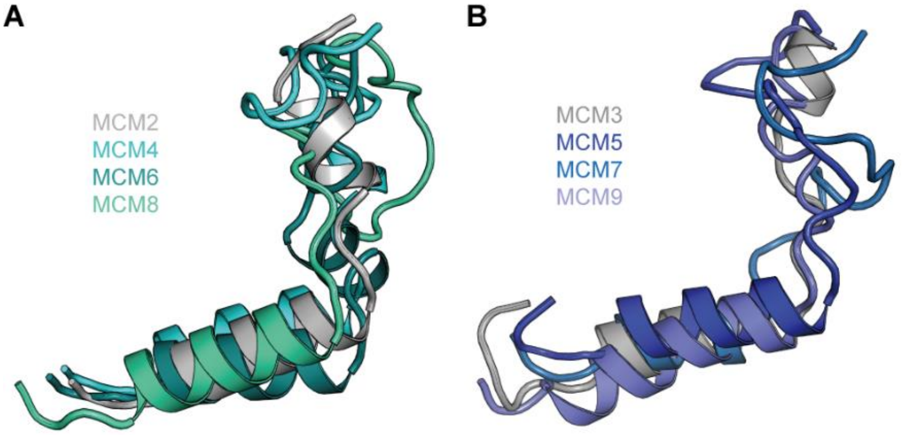


**C**

N-C linker

```
MCM2 NTD- KKDNKVA...............VGELTDEDVKMITSLSKDQQ -CTD 27
MCM4 NTD- KTDAKRLHGLDEEAEQK.......LFSEKRVELLKELSRKPD -CTD 35
MCM6 NTD- ..TNPRFGGKELRDEEQTAESIKNQMTVKEWEKVFEMSQDKN -CTD 40
MCM8 NTD- .....SKGQKTKSSEDGCKHGMLMEFSLKDLYAIQEIQAEEN -CTD 37
```

**D**

N-C linker

```
MCM3 NTD- ...................MSKDAQPSFSAEDI....... -CTD 14
MCM5 NTD- .......DTDGSGRSFAGAVSPQEEEEFRRLAALPNV... -CTD 30
MCM7 NTD- KMNKSEDDESGAGE.....LTREELRQIAEED........ -CTD 27
MCM9 NTD- ......NNEQSSGII....MDEEVQKEFE..DFWEYYKSD -CTD 28
```

**Figure S10. Subgroups of the N-C linkers of MCM2-9**. The N-C linkers of MCM2-9 were divided into two subgroups according to structural and sequence similarity. The N-C linkers of MCM2, MCM4, MCM6 and MCM8 formed group I while the other N-C linkers made up another group II. (**A, B**) structural alignments of the N-C linkers from group I (A) and group II (B). All the structures are from AlphaFold prediction website. (**C, D**) sequence alignments of the N-C linkers of group I (C) and group II (D), respectively. The sequence alignments were performed in ESPript3.0 (*1*). conserved residues are highlighted as red and deletion residues of MCM8 and MCM9 in helicase assay are labelled by red boxes. The length of the N-C linkers of MCM2-9 are listed on the right.

Figure S11

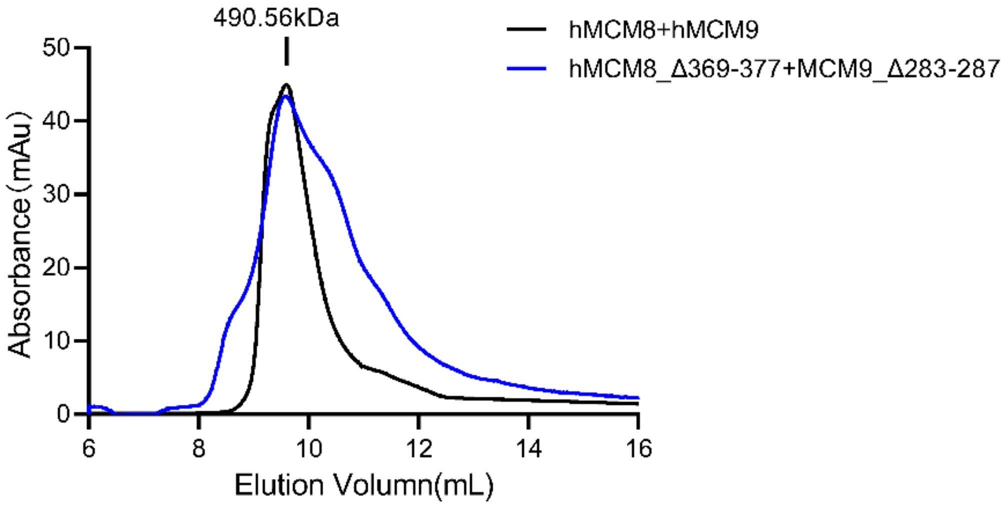


**Figure S11. SEC profiles of WT and N-C linker mutant of MCM8/9 complex.**

Figure S12

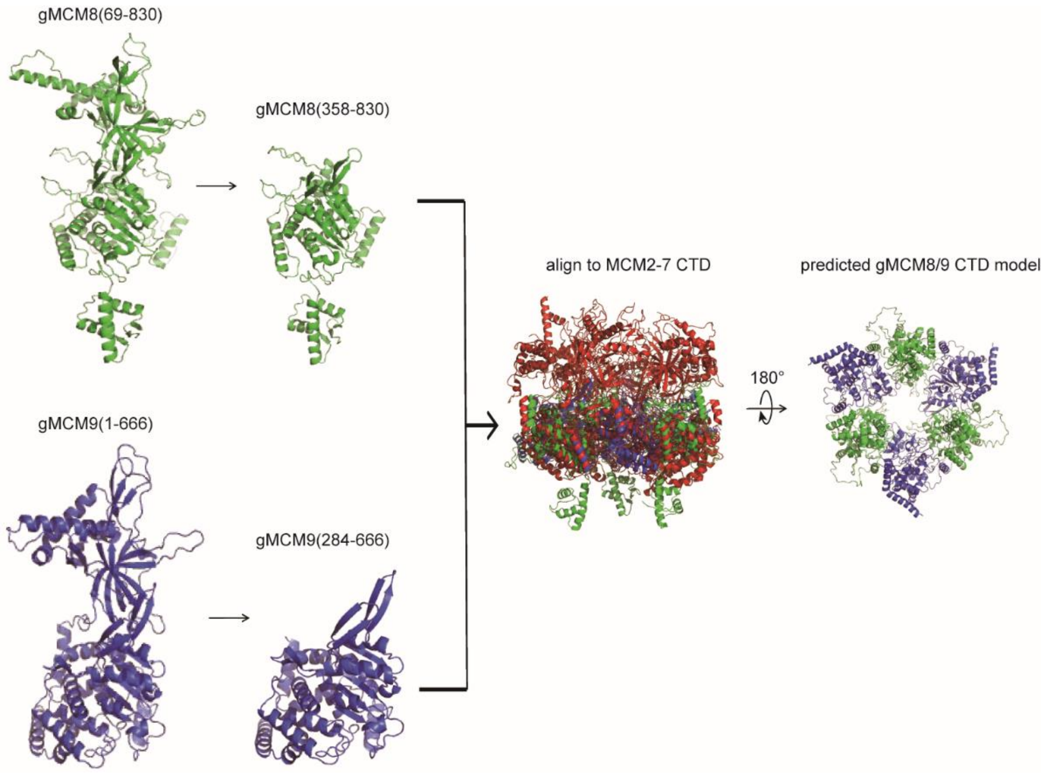


**Figure S12. The predicted model uses as a reference for morphing map of the gMCM8/9 CTD.** The structures of gMCM8 and gMCM9 were predicted using AlphaFold. Their CTD were extracted and aligned to MCM2-7 (PDB: 3JA8) by applying *C3* symmetry.

Figure S13

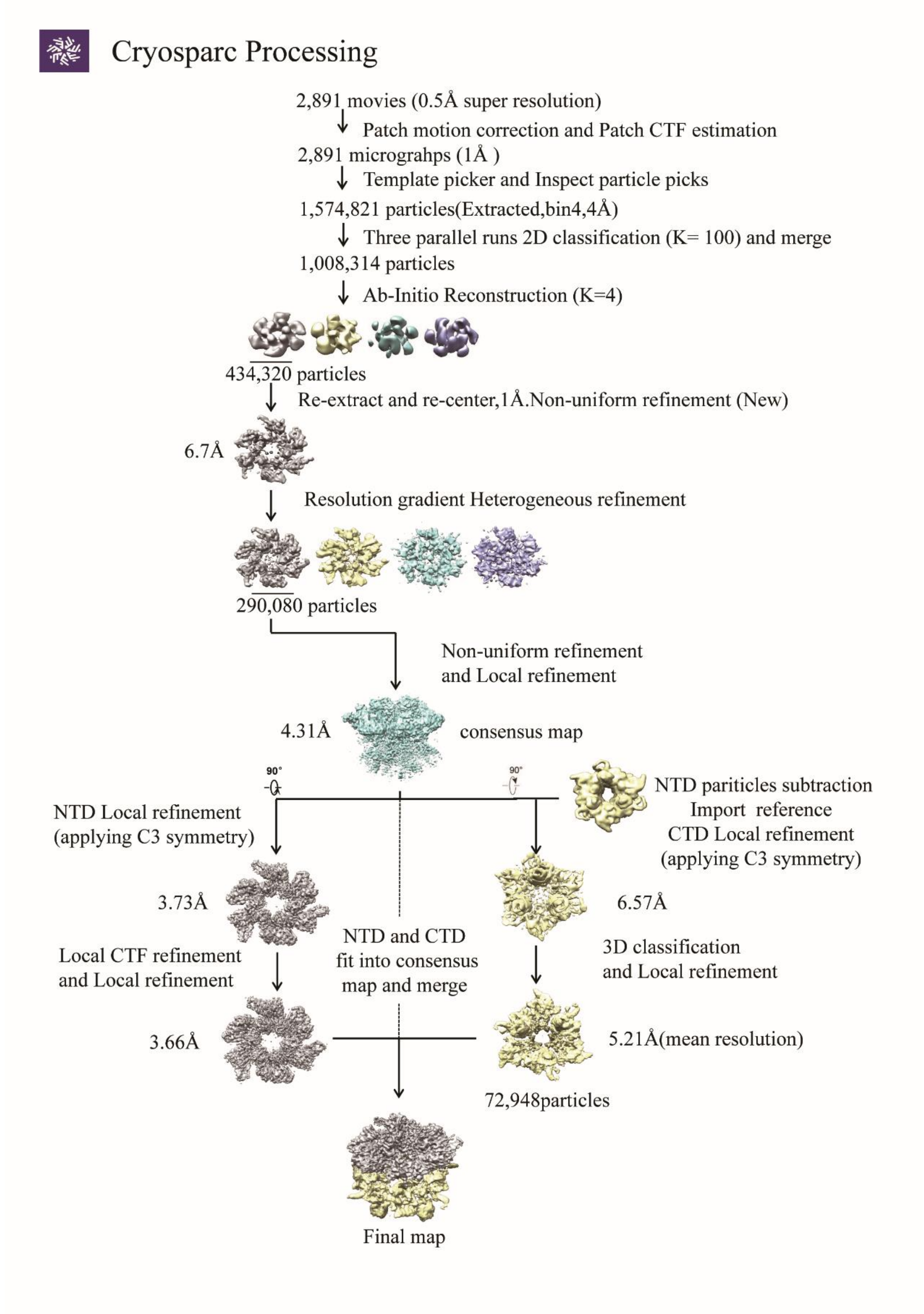


**Figure S13. The image processing and 3D reconstruction steps of the gMCM8/9 complex using cryoSPARC.**

Figure S14

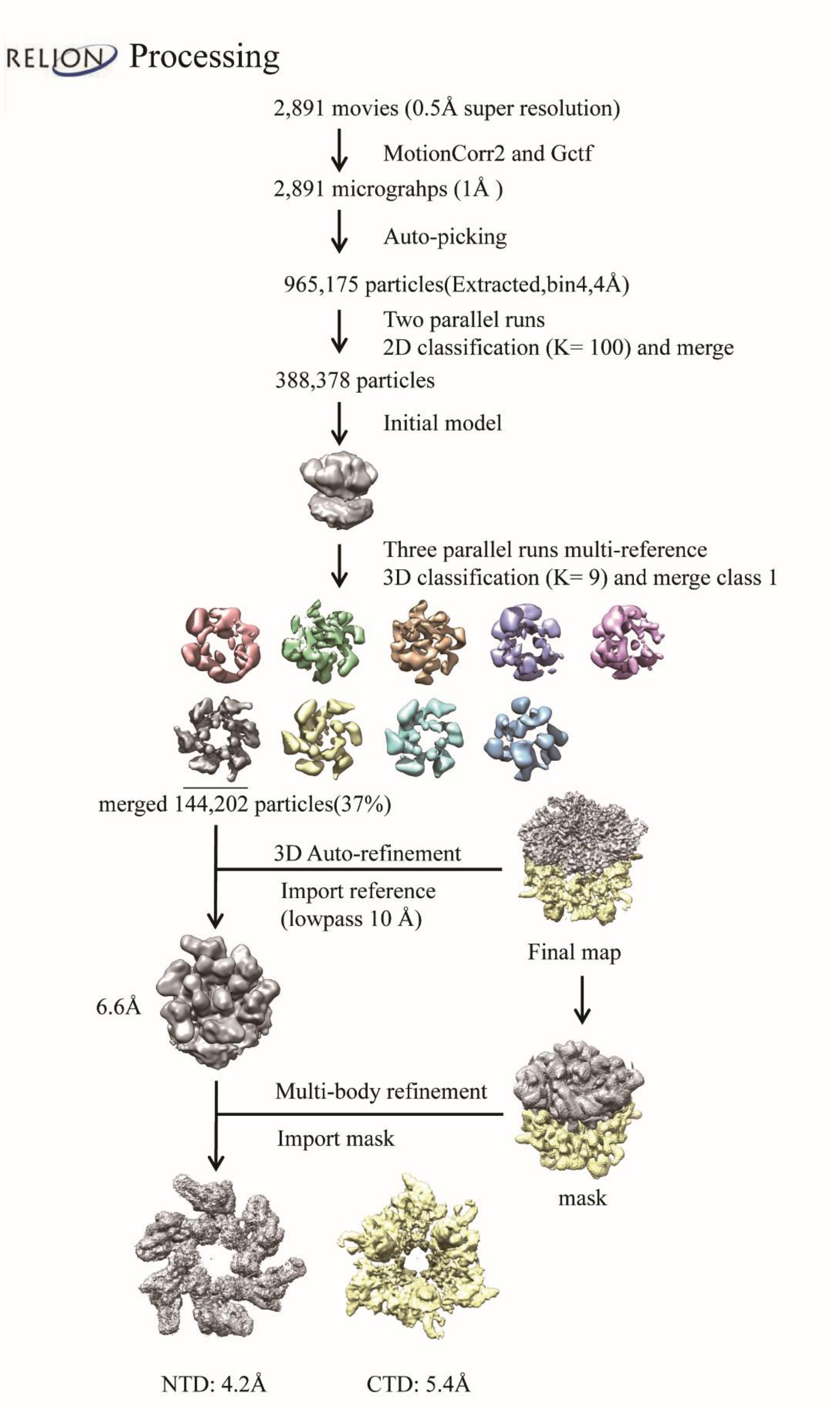


**Figure S14. The image processing and 3D reconstruction steps of the gMCM8/9 complex using RELION-3.1.1.**

Figure S15

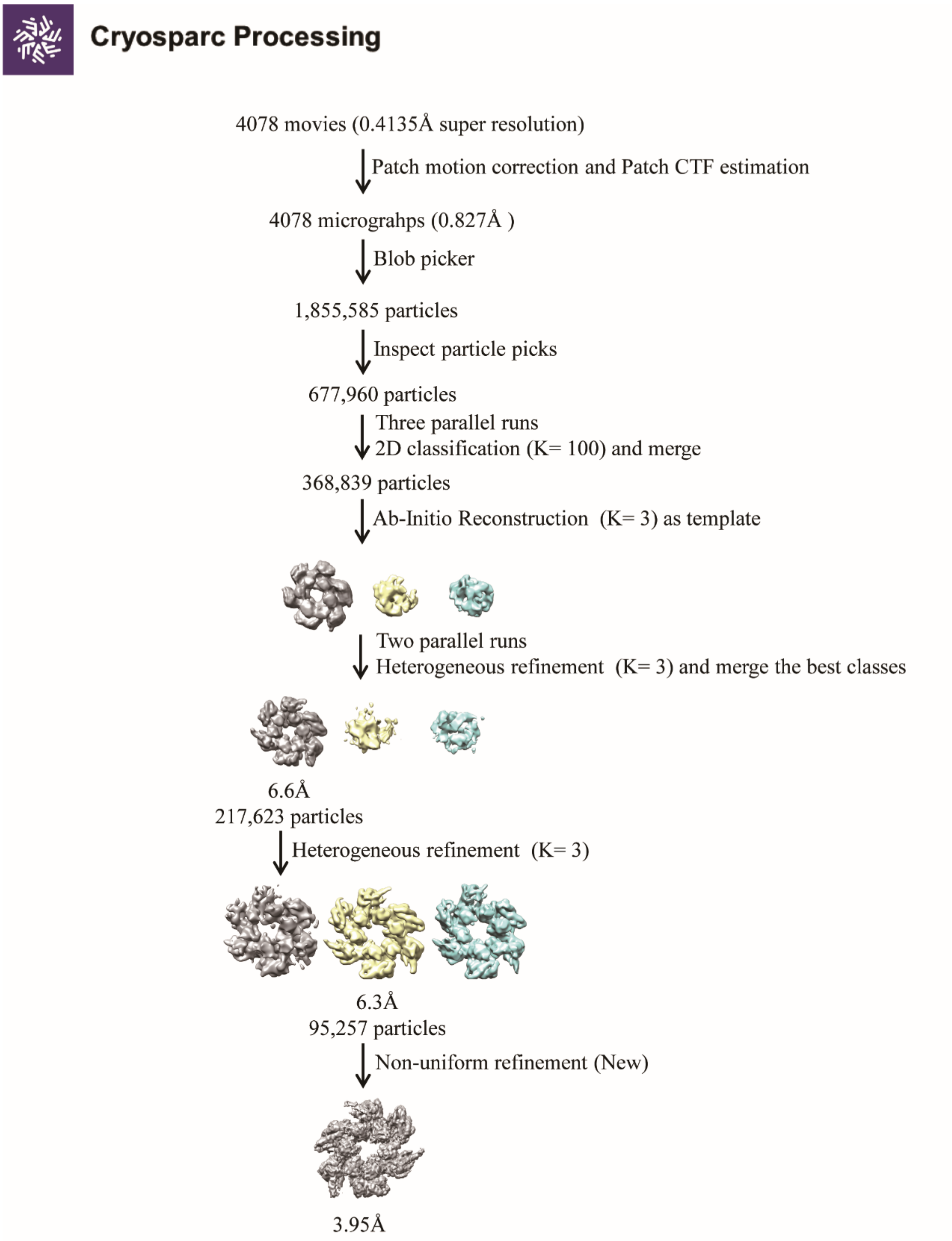


**Figure S15. The image processing and 3D reconstruction steps of the NTD ring of hMCM8/9 Conformation II using cryoSPARC.**
